# Climate Change and Phenological Monitoring of the Indigenous Godello Variety in a Territory of Heroic Viticulture and Biosphere Reserve

**DOI:** 10.1101/2023.12.25.573298

**Authors:** Mª Eva Fernandez-Conde, Dipti Bisarya, Liwayway Perlado Taglinao, J. Antonio Cortiñas

## Abstract

An evaluation of the phenological behaviour of the indigenous Godello variety of vineyard was conducted within the Souto Chao plot in the year 2023. This plot is located in a wine-growing region holding a Designation of Origin (Ribeira Sacra DO) located in the North-West of the Iberian Peninsula. A similar study had been conducted in 2017 and 2018, focusing on the same Godello variety within the identical plot involving the same plant as the current study.

The assessment utilized the BBCH scale, a system proposed by *Biologische Bundesanstalt für Land und Forstwirtschaft, Bundessortenamt und CHemische Industrie*.

The average duration of the vegetative cycle in the initial two years under examination was 167 days (161 and 174 days in 2017 and 2018, respectively), while in 2023 it was 164 days. In these three years studies during 2017, 2018 and 2023, the longest phenological phase was of 79 days when majority of the berries were touching).

The primary objective of this study is to ascertain the nature and extent of changes in the crop’s phenology within the timeframe between the initial two years analysed (2017 and 2018) and the present year, 2023.

## 1. Introduction

Ribeira Sacra is considered one of the greatest examples of what is known as heroic viticulture. This term refers to the terrain conditions which makes it very difficult to work on the vineyards, as it is cultivated on terraces and the slope of the land which are very steep.

It is located in the interior of Galicia (Northwest Spain), it is an area of great scenic wealth, recognised worldwide as a Biosphere Reserve by UNESCO in 2021, by Resolution of 5 October. https://www.boe.es/diario_boe/txt.php?id=BOE-A-2021-16794

The territory includes areas in the south of the province of Lugo and the north of Ourense. The rivers Miño, Sil and Cabe flow through the area, forming meanders and impressive canyons. This is the territory for vine cultivation on high slopes, with landscapes of great biological richness from both the faunal and floral perspectives. The cultivation is located at the places that are very difficult for winegrowers to access, because the orography prevents mechanised cultivation. The information of grapevine phenology over several seasons provides essential information on the characteristics of wine-growing regions, allowing cultural practices and phytosanitary treatments to be better planned. The phenological studies are very important because they are closely related and easily affected by global warming and climate change. The characterisation of its possible behaviour could serve as a reference for the definition of strategies in crop planning, especially for vineyards due to their great importance in this area.

The information about and major changes in the development cycle of the plants due to rising temperatures would lead to better understanding of the management of the crop. Addressing the impact of potential climate change on vine phenology will allow the wine industry in future to define some strategies that could be useful for further vineyard planning, such as shifting the cultivation of grapes to more northerly areas that have comparatively milder temperatures. One of the examples is the major expansion seen in United Kingdom (UK), where the area under vines has increased by about 150% in recent years as a result of global warming and climate change [1, 2].

The study focuses on a Designation of Origin (DO) of enormous importance in the production of various wines, with great character and wide acceptance in international markets. The initial studies on phenology and phytopathology in Ribeira Sacra vineyards were conducted in 2017 [3, 4].

Recent data could help us to understand the effects of climate change on vine cultivation, a plant that is very sensitive to climatic variations, and to know its response to the different scenarios produced [5].

The study of the phenological phases of the vine reflects how the development and growth of vegetative and fruiting organs takes place, and there is a close relationship with climatic and cultural conditions [1].

It has been observed that the harvesting dates have been brought forward considerably in the period between the end of the 20th century and the beginning of the 21st century to avoid the accumulation of excess alcohol content and also to preserve the freshness of the wines, but this in turn brings another major problem; the skins and seeds, which gives the wine important organoleptic properties such as astringency, colour and aromas, do not ripen at the same time and rate as the pulp and develop slowly. In white wines, this results into the wines that is heavier and harder on the palate, less agile and also difficult to preserve. However, this problem is not universal for all grape varieties and different regions. For example, Galician white wines, such as the Godello variety, where viticulture is adapted to cooler areas, have become fresher and more aromatic, therefore have gained a greater market share [6].

Climate change has become one of the world’s major concerns for wine production, as vine cultivation is very sensitive to changes in climatic conditions, especially changes in temperature and solar radiation. These factors are crucial for the ripening of the fruit, which means that harvests are being shifted little earlier, with the aforementioned risks of altering the chemical composition of the grape and the organoleptic characteristics of the product obtained [7, 8]. In addition to the changes in the chemical composition of the wine, the increase in temperature causes pests and plants to react to these changes. Temperatures are not only the result of warmer summer days, but also of fewer cold days and less frost, which favours the growth of fungi and pests, altering the interaction of the disease triangle (host-pathogen-environment) and thus ultimately reducing the crop production [9, 10].

## 2. Materials and Methods

### 2.1. Location and climatic characteristics of the study area

The study was conducted in one vineyard in Ribeira Sacra (Souto Chao) during the year 2023, and previously the studies were conducted in years 2017 and 2018 (Figure 1).

**Figure 1.**
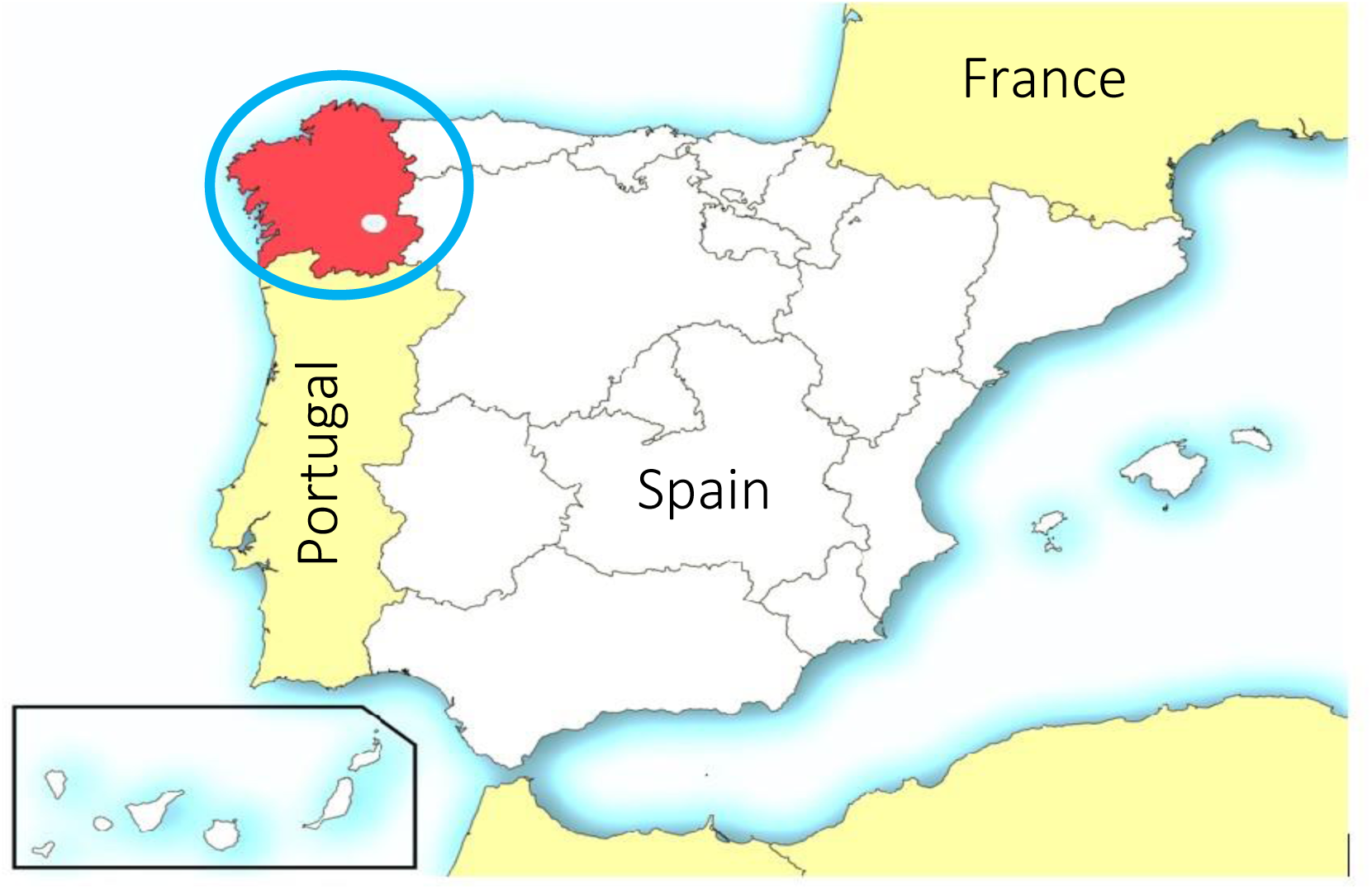
Location of the Ribeira Sacra area in the North-West of the Iberian Peninsula (Spain)

Ribeira Sacra comprises 2,500 ha of vineyards. The orography of the terrain requires the vines to be planted on very steep slopes, which the *Centro di Ricerca, Studi, Salvaguardia, Coordinamento e Valorizzazione per la Viticoltura di Montagna*, CERVIM, describes as “heroic viticulture”. http://www.cervim.org/

The Souto Chao vineyard is located at 438 m.a.s.l. on the lower terraces along the banks of the River Sil, following the contours and with slopes of up to 80% (42°24′27.67″ N, 7° 28′20.06″ W). The soils are shallow and of low fertility, which limits the growth of the vine shoots [11].

During the warmest month, temperatures on the terraces closest to the river (Souto Chao) can be up to one and a half degrees higher than the average for Ribeira Sacra as a whole. The river has a significant thermoregulatory effect, especially when combined with the favourable South-Southeast exposure of the vineyards. On the other hand, in the higher parts, especially in the areas where orientation is not so favourable for viticulture, there is a negative temperature variation of similar magnitude [12].

The Godello variety has small to medium sized grapes, conical in shape, with fairly uniform berries of medium to high compactness, with a medium sized stalk and little lignification at the base. The leaf is medium to large, pentagonal and has five lobes [13].

The weather data was collected and the average maximum and minimum temperature, relative humidity, hours of sunshine and rainfall are presented in Table 1. The data was collected from the station located near to vineyard study area, which belongs to the Meteogalicia weather service. https://www.meteogalicia.gal.

**Table 1.**
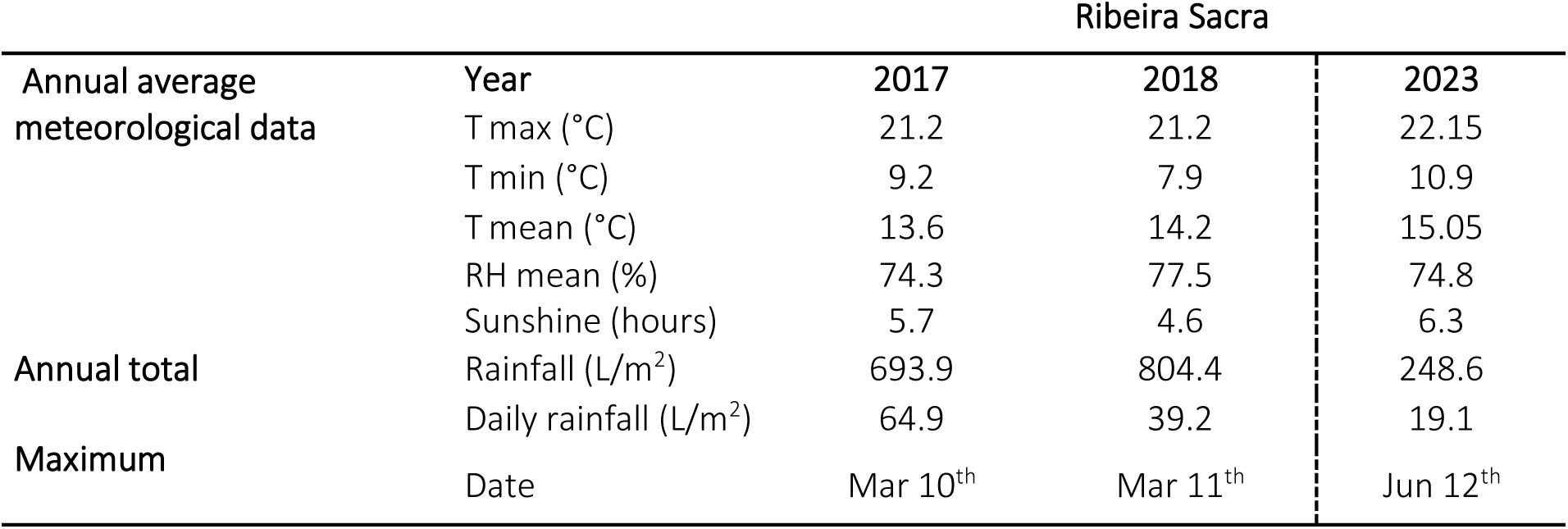
Weather data of the study area during de crop season (2017, 2018 and 2023)

### 2.2. Phenological protocol and Fieldwork

The initial phenological studies in the Souto Chao plot were carried out in 2017 and 2018. After 5 years, the phenology was studied again on the plants of same variety from the same plot. Although the time interval is not very long, we tried to obtain some important data on how climate change affected the vine cultivation in this area of particular orography and terraced cultivation systems on the slopes of the river.

We studied the phenological stages of the Godello cultivar according to the scale adopted by the BBCH, which is the standardised scale by Lorenz [14] (Table 2). The phenological observations were applied to randomly selected 10 vines distributed throughout the plot. The study was carried out during the active grapevine cycle, from the 1st of March 2023 until the grape harvest in last week of August 2023. During the sampling period, the vineyard was visited weekly, except during the flowering period, when the number of visits was increased to twice a week. In order to determine the phenological calendar of the vines, the beginning of each stage was considered to be when 50% of the plants under study had reached that particular stage. From the BBCH scale, a total of 18 phenological phases belonging to the six main stages were selected for observation.

**Table 2.** Stages and Phases of the BBCH-Skala considered in this study.

| STAGES (St) | PHASES (Ph) |
| --- | --- |
| 0: Sprouting | 03 - End of bud swelling: buds swollen, but not green |
|  | 05 - "Wool stage": brown wool clearly visible |
|  | 09 - Bud burst: green shoot tips clearly visible |
| 1: Leaf development | 11 - First leaf unfolded and spread away from shoot |
|  | 13 - 3rd leaves unfolded (continuous) |
|  | 15 - 5th leaves unfolded |
| 5: Inflorescence emerge | 53 - Inflorescences clearly visible |
|  | 55 - Inflorescences swelling, flowers closely pressed together |
|  | 57 - Inflorescences fully developed; flowers separating |
| 6: Flowering | 61 - Beginning of flowering: 10% of flowerhoods fallen |
|  | 65 - Full flowering: 50% of flowerhoods fallen |
|  | 69 - End of flowering |
| 7: Development of fruits | 71 - Fruit set: young fruits begin to swell, remains of flowers lost |
|  | 75 - Berries pea-sized, bunches hang |
|  | 79 - Majority of berries touching |
| 8: Ripening of berries | 81 - Beginning of ripening: berries begin to develop variety-specific colour |
|  | 85 - Softening of berries |
|  | 89 - Berries ripe for harvest |

### 2.3. Thermal heat requirements

Heat requirements were calculated according to the criterion proposed by Galan [15], which took into account the daily sum of maximum temperatures (Growing Degrees-Days, GDD) from the end of the cold period to the beginning of each phenological phase.GDD °C =∑ Ti_max_

Ti _max_= *daily maximum temperatures in a number of days i*.

The F-test of statistical analysis determines whether the variability between the means of the groups is greater than the variability of the observations within the groups. If this ratio is sufficiently large, it can be concluded that not all means are equal. In this study it gives a relatively high value, due to the fact that it is a one-year sample where the sample size was not too large (Table 3).

**Table 3.**
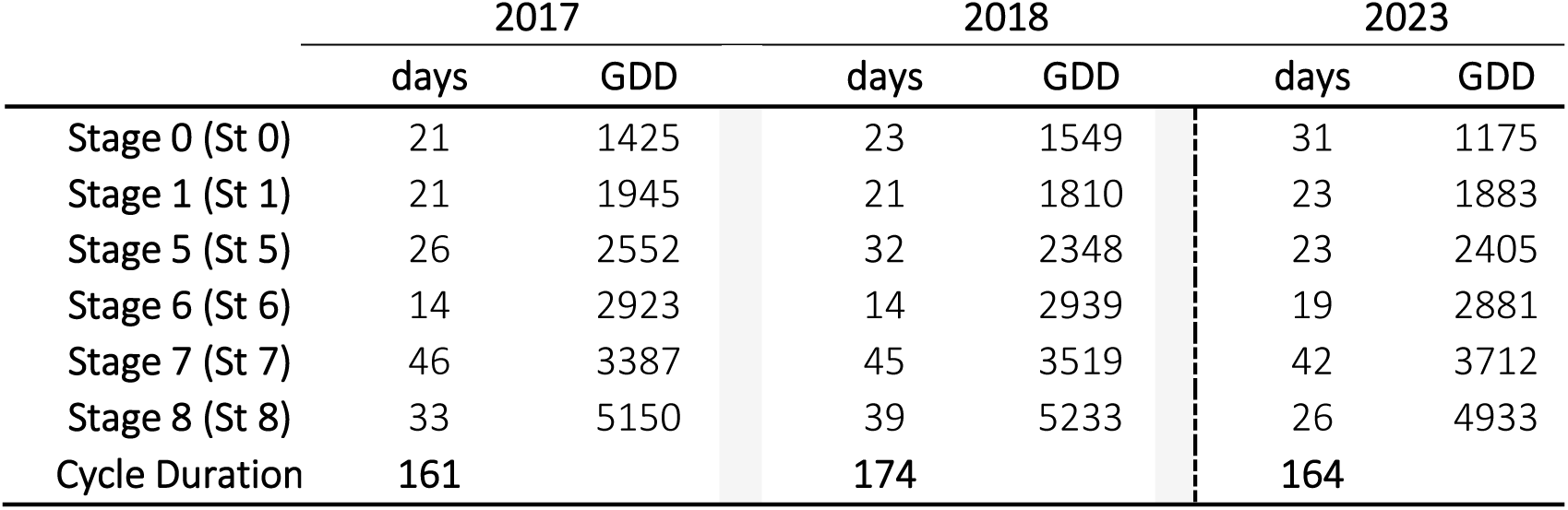
Duration of the stages and thermal requirements (growing degrees-days, GDD)

|  | 2017 |  | 2018 |  | 2023 |  |
| --- | --- | --- | --- | --- | --- | --- |
|  | days | GDD | days | GDD | days | GDD |
| Stage 0 (St 0) | 21 | 1425 | 23 | 1549 | 31 | 1175 |
| Stage 1 (St 1) | 21 | 1945 | 21 | 1810 | 23 | 1883 |
| Stage 5 (St 5) | 26 | 2552 | 32 | 2348 | 23 | 2405 |
| Stage 6 (St 6) | 14 | 2923 | 14 | 2939 | 19 | 2881 |
| Stage 7 (St 7) | 46 | 3387 | 45 | 3519 | 42 | 3712 |
| Stage 8 (St 8) | 33 | 5150 | 39 | 5233 | 26 | 4933 |
| Cycle Duration | 161 |  | 174 |  | 164 |  |

The optimum temperatures requirements for this crop are different at different stages of development, so it is necessary to determine these conditions.

### 2.4. Statistical Analysis

Two statistical tests were used to determine possible differences in the length of the stage variation, within and between groups: two-way analysis of variance (ANOVA) and Tukey’s honest significant differences (Tukey’s HSD).

Test ANOVA: by stage-Godello variety (Df: degrees of freedom; Sum Sq: Sum of square; Mean Sq: mean square; plevel=0.1) and Tukey HSD between Stages (Lower and Higher are the ends of the confidence interval; plevel=0.05). TUKEY HSD/KRAMER Qtest: between groups (mean; std err; q-stat; lower; upper, p-value; mean-crit).

## 3. Results

### 3.1. Phenological behaviour of the Godello variety

The vegetative growth period of the Godello variety was calculated from the end of bud swelling, when the buds were swollen but not green (phase 03), stage 0: sprouting, to the berries ripe for harvesting (phase 89), stage 8: Ripening of the berries. During the first study in 2017, the vegetative cycle of the vine was from March to August, while in 2018 the vegetative cycle was from April to September.

In the year 2017, which was the first year of study, the tagging was done and records have been kept of the plants in which harvesting was done for the first time in August 2017. In this wine-growing region, harvesting is usually carried out in the month of September. In this present study of 2023, the grape growing season was again from March to August, i.e., for the second time the grapes were harvested again in August 2023, specifically in the second fortnight.

The drought and high spring temperatures are resulting into early harvesting of the crop in the areas such as Ribeira Sacra, which is threatening and leading to reduced production. The variety Godello is easily affected and is also sensitive to this new situation. Due to the fact that the high temperatures arrive early and accelerate the ripening process, vine growers and wine makers are indicating that the harvesting must begin four to five days earlier each year, which also depends on the variety, with the white variety i.e., Godello which is being easily affected. In the three years of our study, the longest phenological phase (Ph) was 79 (majority of berries touching), with a duration of 34, 27 and 20 days in 2017, 2018 and 2023 respectively.

The shortest phase in 2017 was Ph 61 (Beginning of flowering: 10 % of flowerhoods fallen) with a duration of 3 days. In 2018, Ph 11 (First leaf unfolded and spread away from shoot); Ph 65 (Full flowering: 50 % of flowerhoods fallen) and Ph 69 (End of flowering), with a duration of 3 days each. In the present study carried out in 2023, the shortest phase was Ph 55 (Inflorescences swelling, flowers closely pressed together) with a duration of 5 days. The differences between the longest and shortest phases in 2017, 2018 and 2023 were 31, 24 and 15 days respectively. It follows that in the latter year there was less contrast in terms of the duration of the different phenological phases.

The difference between the longest and the shortest stage (St) in 2017, 2018 and 2023 was 32, 31 and 23 days, respectively (St 7: development of fruits and St 6: flowering). In terms of the phenological phases, the phenological stages in the last year of the study showed a smaller difference in duration between them (Table 4).

**Table 4.** Average dates of the beginning of each phenological phase and duration in the different years for the Godello variety.

| Stages | Phases | Start Date |  | Duration of phase |  |  | Start date | Duration of phase |
| --- | --- | --- | --- | --- | --- | --- | --- | --- |
|  |  | 2017 | 2018 | 2017 | 2018 | Mean | 2023 |  |
| 0 | 3 | 18-Mar | 3-Apr | 7 | 12 | 10 | 17-Mar | 8 |
|  | 5 | 25-Mar | 15-Apr | 8 | 7 | 8 | 25-Mar | 7 |
|  | 9 | 2-Apr | 22-Apr | 6 | 4 | 5 | 31-Mar | 16 |
| 1 | 11 | 8-Apr | 28-Apr | 7 | 3 | 5 | 16-Apr | 8 |
|  | 13 | 15-Apr | 1-May | 7 | 9 | 8 | 24-Apr | 7 |
|  | 15 | 22-Apr | 10-May | 7 | 9 | 8 | 1-May | 8 |
| 5 | 53 | 29-Apr | 19-May | 8 | 4 | 6 | 9-May | 10 |
|  | 55 | 6-May | 23-May | 14 | 10 | 12 | 19-May | 5 |
|  | 57 | 20-May | 2-Jun | 4 | 18 | 11 | 24-May | 8 |
| 6 | 61 | 24-May | 20-Jun | 3 | 8 | 6 | 1-Jun | 6 |
|  | 65 | 27-May | 28-Jun | 7 | 3 | 5 | 7-Jun | 6 |
|  | 69 | 3-Jun | 1-Jul | 4 | 3 | 4 | 13-Jun | 7 |
| 7 | 71 | 7-Jun | 4-Jul | 4 | 4 | 4 | 20-Jun | 7 |
|  | 75 | 11-Jun | 8-Jul | 8 | 14 | 11 | 27-Jun | 15 |
|  | 79 | 19-Jun | 22-Jul | 34 | 27 | 31 | 12-Jul | 20 |
| 8 | 81 | 22-Jul | 18-Aug | 6 | 7 | 7 | 1-Aug | 8 |
|  | 85 | 28-Jul | 25-Aug | 22 | 22 | 22 | 10-Aug | 12 |
|  | 89 | 19-Aug | 16-Sep | 5 | 10 | 8 | 22-Aug | 6 |

The average length of the growing season in the two years was 167 days (161 and 174 days in 2017 and 2018, respectively), while in 2023 it was 164 days. Considering 2017 as exceptional, where the harvest was brought forward to August for the first time, the year 2023 confirms once again this exceptionality; the harvest was again in August.

All the evidences, point us to change the trend to a relatively short period of time. Furthermore, phenological studies in the future should confirm this working hypothesis.

The standard deviation of the average start date of the various phenological stages exhibited lower values for the period 2017/2023 (6 compared to 16 for 2017/2018 and 10 for 2018/2023). This result indicates a lower variability of the start dates of the phenological stages from one period to another. Regarding the mean values of the duration of each phenological stage, the values are similar for the three periods studied (value of 3), indicating that there are no significant differences. The lowest variability in the average start date of the different phenological stages for the period 2017/2023 corresponds to the only two years recorded so far, where sprouting and harvest took place from March to August. (Figure 2a; 2b).

**Figure 2a.**
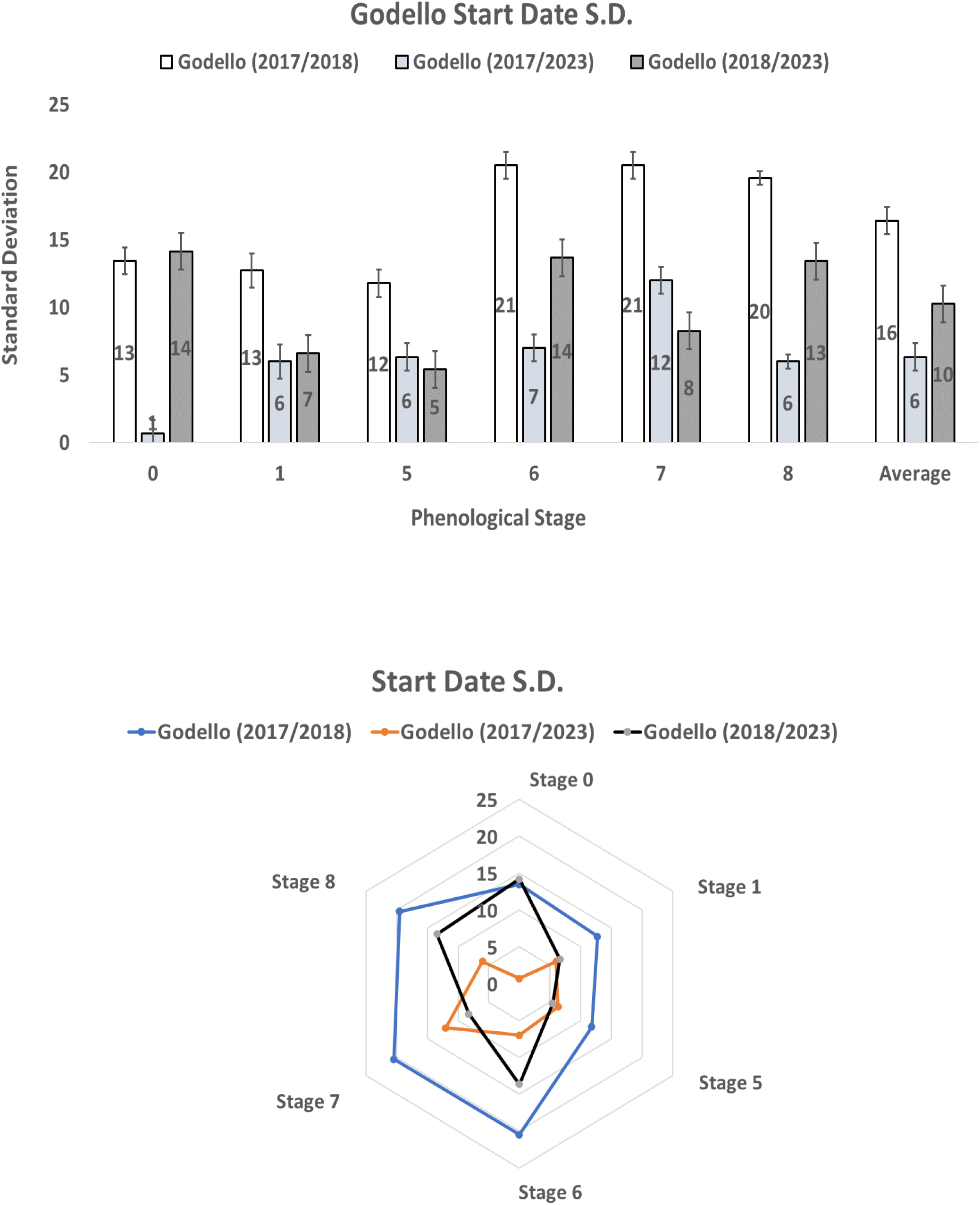
Start date of the Standard Deviation (S. D.) for each phenological stage with the average values (Error bars included in the first graph)

**Figure 2b.**
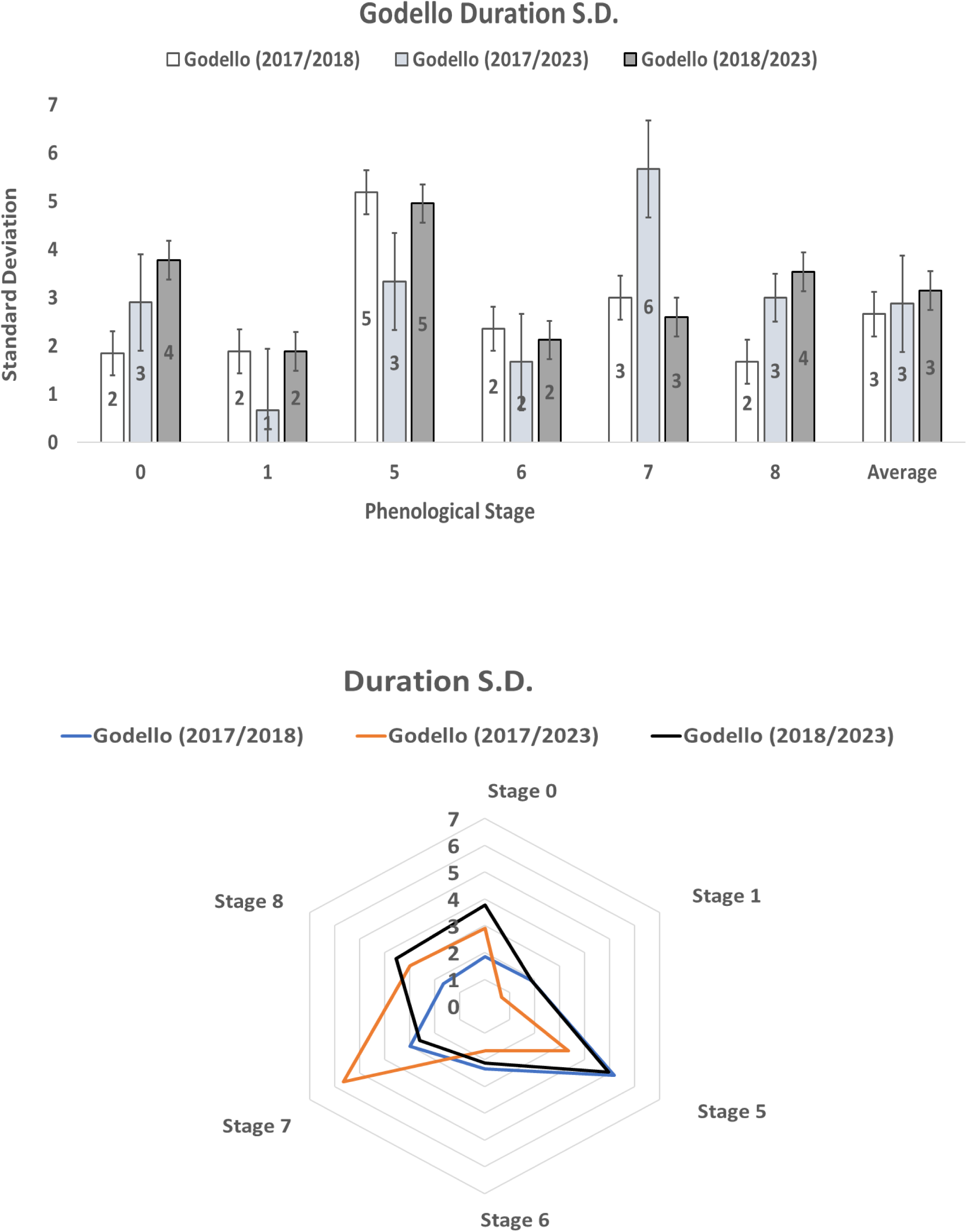
Duration of the Standard Deviation (S. D.) for each phenological stage with the average values. (Error bars included in the first graph)

### 3.2. Analysis of meteorological parameters

The phenological phases included in main stage 0 (St 0: Sprouting) take place this year from mid-March to the end of March. During this phase, the lowest average maximum and minimum temperatures (17.2°C and 6.6°C, respectively) were recorded. Stage 1 (St 1: Leaf development) takes place from mid-April to early May. From then until the beginning of June, stage 5 (St 5: Inflorescence emerge) took place.

The next stage 6 (St 6: Flowering) took place from the beginning to the middle of June. The highest rainfall value (19.1 mm of rainfall) was recorded at this stage, namely on 12 June. Stage 7 (St 7: Development of fruits), runs from June to the beginning of August. Finally, the last phenological stage of the vegetative cycle of the vine 8 (St 8: Ripening of berries), occurred between the beginning of August and the end of the same month, recording the highest average maximum, minimum and average temperatures (29.9°C, 14.8°C and 21.7°C, respectively) (Figure 3).

**Figure 3.**
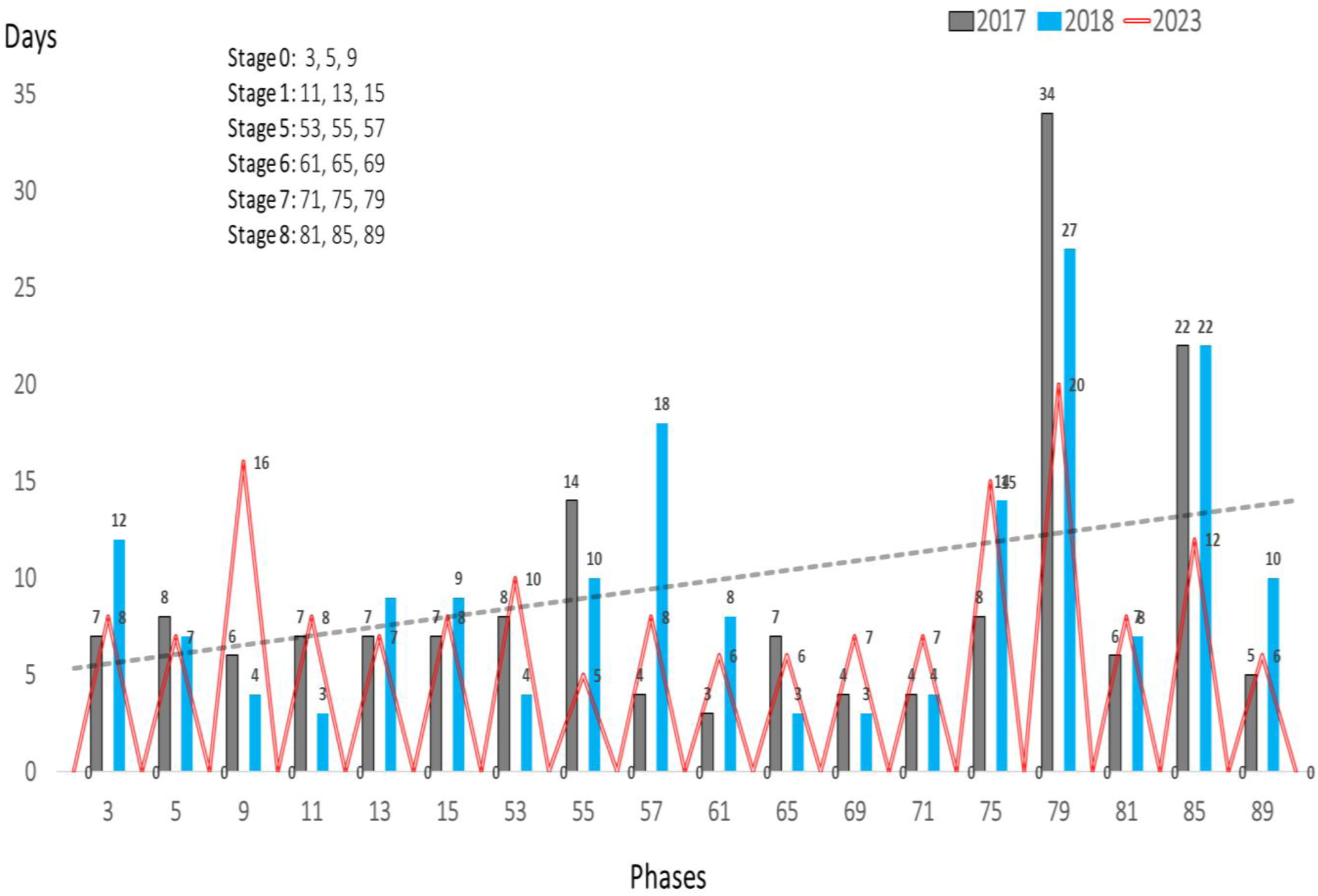
Length of each phenological phase in the different years of study

The accumulated heat units for berry ripening (St 8) reached the levels of around 5000 GDD. Thus, the thermal requirements needed to overcome the different phenological stages ranged from 1175 GDD for stage 0 (St 0) to 4933 for stage 8 (St 8). (Table 3).

Heat requirements were determined by calculating the accumulated sum of daily maximum temperatures (in growing degree days, GDD) from the conclusion of the cold period until the onset of each phenological phase for the Godello Variety (*Vitis vinifera* L.) grown in the Souto Chao plot on the slopes of the Sil River, where the particular orography of this territory, with slopes of up to 80%, as well as the proximity to the river, determine the conditions for vine cultivation.

In these areas it is essential to conserve the scarce soil moisture and to reduce erosion processes. One of the techniques used by winegrowers is mulching, the main objective of which is to prevent the proliferation of weeds. Non-living mulches have a number of advantages. These include improved soil structure, disease and pest control, crop quality improvement, which reaps financial benefits. The primary benefits, however, are associated with weed control. Weed suppression can result in significant long-term labour and herbicide savings. It is critical to select the appropriate mulch for the situation. The effectiveness of the mulch will vary depending on the prevailing weed problems and the surrounding environment [16, 17] (Figure 4).

**Figure 4.**
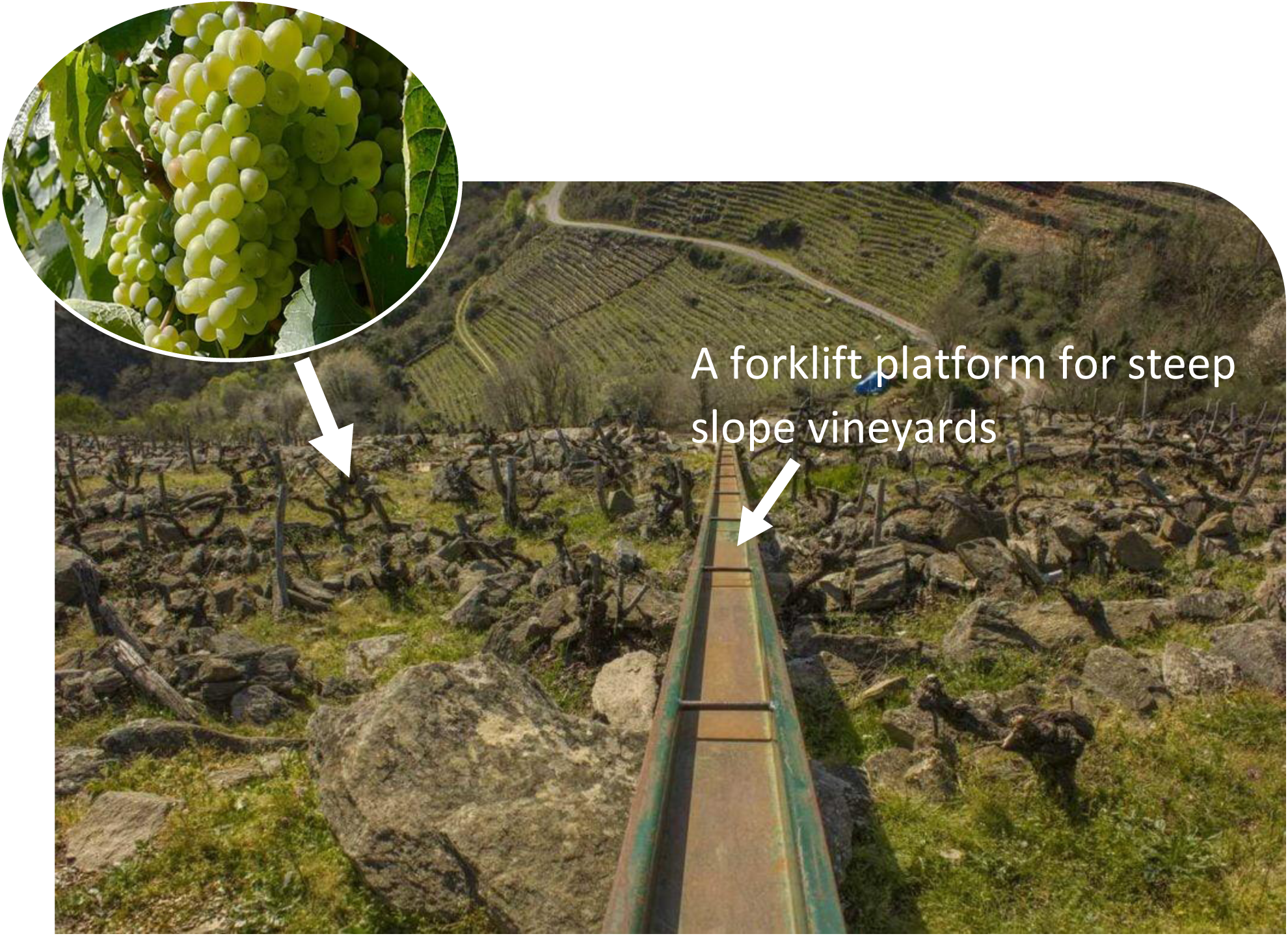
Vineyards in Ribeira Sacra. Godello variety in the Souto Chao plot

### 3.3. Statistical Analysis

The ANOVA test results show that no significant differences were found between the samples analysed in terms of the duration of each stage and phenological stage in the different years of the study.

The calculated *F* value was lower than the *critical F* value. The *alpha* value (0.05) is also lower than the *p-value* (0.93835). In the samples analysed, the value found is 0.06370, which is lower than the value of 3.17879.

This result has also been confirmed by the Tukey HSD/KRAMER Q test where the *alpha* value 0.05, is lower than the *p-value* calculated between the different groups with values of: 0.93804; 0.9965 and 0.96284 (Table 5).

**Table 5.**
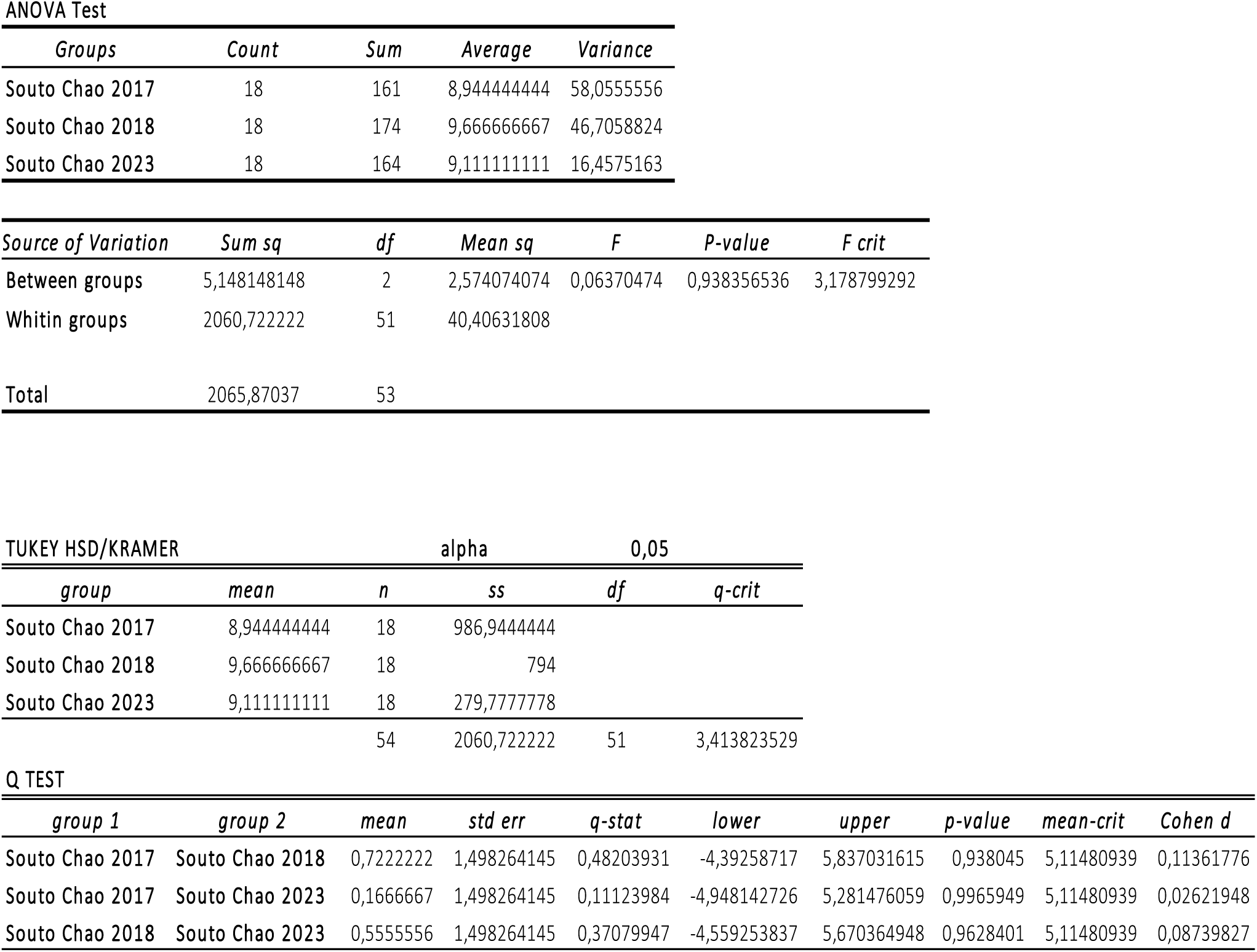
Statistical analysis in the different years. The values of the Test ANOVA were also shown and calculated: Df=degrees of freedom; Sum Sq=sum of square; Mean Sq=mean square; F; P-value and F crit. The TUKEY HSD/KRAMER was also calculated between the different groups (mean; std err; q-stat; lower; upper; p-value; mean-crit and Cohen d)

ANOVA Test
| *Groups* | *Count* | *Sum* | *Average* | *Variance* |
| --- | --- | --- | --- | --- |
| Souto Chao 2017 | 18 | 161 | 8,944444444 | 58,0555556 |
| Souto Chao 2018 | 18 | 174 | 9,666666667 | 46,7058824 |
| Souto Chao 2023 | 18 | 164 | 9,111111111 | 16,4575163 |

| *Source of Variation* | *Sum sq* | *df* | *Mean sq* | *F* | *P-value* | *F crit* |
| --- | --- | --- | --- | --- | --- | --- |
| Between groups | 5,148148148 | 2 | 2,574074074 | 0,06370474 | 0,938356536 | 3,178799292 |
| Whitin groups | 2060,722222 | 51 | 40,40631808 |  |  |  |
| Total | 2065,87037 | 53 |  |  |  |  |

TUKEY HSD/KRAMER alpha 0,05
| *group* | *mean* | *n* | *ss* | *df* | *q-crit* |
| --- | --- | --- | --- | --- | --- |
| Souto Chao 2017 | 8,944444444 | 18 | 986,9444444 |  |  |
| Souto Chao 2018 | 9,666666667 | 18 | 794 |  |  |
| Souto Chao 2023 | 9,111111111 | 18 | 279,7777778 |  |  |
|  |  | 54 | 2060,722222 | 51 | 3,413823529 |

**Table 5** Statistical analysis in the different years.
| *group 1* | *group 2* | *mean* | *std err* | *q-stat* | *lower* | *upper* | *p-value* | *mean-crit* | *Cohen d* |
| --- | --- | --- | --- | --- | --- | --- | --- | --- | --- |
| Souto Chao 2017 | Souto Chao 2018 | 0,7222222 | 1,498264145 | 0,48203931 | -4,39258717 | 5,837031615 | 0,938045 | 5,11480939 | 0,11361776 |
| Souto Chao 2017 | Souto Chao 2023 | 0,1666667 | 1,498264145 | 0,11123984 | -4,948142726 | 5,281476059 | 0,9965949 | 5,11480939 | 0,02621948 |
| Souto Chao 2018 | Souto Chao 2023 | 0,5555556 | 1,498264145 | 0,37079947 | -4,559253837 | 5,670364948 | 0,9628401 | 5,11480939 | 0,08739827 |

## 4. Discussion

The active vegetative period of the vine (considered from bud burst to harvest) was longer in 2018, while it was similar in the other two years (2017 and 2023). However, the length of the Godello cycle was 161, 174 and 164 days in 2017, 2018 and 2023 respectively.

Previous studies carried out in this wine region on Mencía, the predominant variety, reported very similar results to those obtained here for Godello [8]. The fact that 2018 shows a significant difference compared to the other two years of study can be attributed to the higher minimum temperatures recorded throughout the vegetative cycle during that year, which leads to a longer development time of the different phenological phases. The lower minimum temperatures in 2018 would explain the longer Godello cycle in the Souto Chao vineyard. A study carried out by Queijeiro [18] in the Ribeiro area showed significantly higher values than those observed for Godello in Ribeira Sacra.

The longest cycle length was observed in parallel studies carried out in the same wine region with other red varieties, such as Brancellao and Merenzao, where a vegetative cycle length was reported to be of 200 days [3].

In other Spanish wine regions (Rioja DO), red varieties such as Tempranillo reported a vegetative cycle length very similar to Godello in Ribeira Sacra in 2018, which was an average of 174 days. In the same Rioja DO, minority red varieties such as Maturana tinta, Maturana tinta Navarrete and Monastel de Rioja reported vegetative cycle lengths of 158, 163 and 173 days, respectively [19]. The phenological cycles of some Turkish varieties have been reported to range from 120 to 176 days, with Trakya Ilkeren and Yalova being the earliest and Favli the latest [20].

The present study showed that the year 2023 was the driest as compared to other years of study in Ribeira Sacra and consequently, the beginning of the berries ripe for harvest (Ph 89) was early, which occurred in mid-August 2023.

Regarding thermal heat requirements, the daily sum of maximum temperatures fluctuated from the sprouting phase (St 0: Sprouting) to berry ripening (St 8: Ripening of berries) stage, in values between 1175 GDD and 4933 GDD. In studies conducted in this geographical area, heat accumulation values recorded for the same variety and in the same plot reported similar values for sprouting (St 0: Sprouting), of 1425 and 1549 GDD, in the years 2017 and 2018, respectively. Values for berry ripening (St 8: Ripening of berries) were 5150 GDD in 2017 and 5233 GDD in 2018, values were in line with those obtained in this present year 2023 of study. The thermal requirements needed to induce flowering (St 6: Flowering) were 2881 GDD, also similar to those obtained in previous studies, which ranged from 2923 to 2939 GDD [3].

This suggest that we must consider the date of the beginning of the flowering stage (St 6: Flowering) because is very important in determining the harvest date, as there is a constant number of accumulated temperature units between these events [19].

Finally, regarding the accumulated thermal units for berry ripening (St 8: Ripening of berries) the values achieved in 2023 were slightly lower than those recorded in 2017 and 2018 in the same plot. In the other viticultural regions also for the red grape cultivars such as Sangiovese and Cabernet Franc, the heat requirements of these varieties during the period between budbreak and flowering varied between 96 and 271 GDD, with the Cabernet Franc variety requiring the highest number of days and the highest sum of heat during the period [21]. These values were very distant from those achieved in the Souto Chao plot for Godello. Therefore, the knowledge of the different phenological stages of the vineyard and their identification is very important for cultural practices and the use of phytosanitary products to control pathogens [2, 22].

Jones and Davis [23] also pointed out that vine development occurs as a direct effect of climate and can be described by phenological events, and stated that the phenology of a crop is important in determining the ability of an area or region to produce crops within the scheme of its climatic regime. The phenological data obtained through a descriptive analysis of the means and standard deviations for the different stages of the vine, from Sprouting (St 0) to Ripening of fruits (St 8), show different values from those obtained in other similar studies for varieties such as Perlon, Red Globe, A. Lavalleé, M. Palieri, Sultanina, etc., but the highest standard deviation was found to be for the variety Perlon. However, the highest standard deviation also coincides with our study in terms of the stages of Flowering (St 6) and Ripening of fruits (St 8) [24].

## 5. Conclusions

It can be concluded that the active vegetative cycle of the vine was generally longer in the year 2018. However, the significant differences were observed between the beginning and end of the different phenological phases for the three years. Both in 2017 and in the present year of study i.e., 2023, the beginning and the end of the vegetative cycle took place between March and August, while in 2018 it was from April to September. The phenological behaviour of the Godello variety is linked to climatic conditions, with temperature being the main meteorological parameter explaining the inter-annual differences in the vineyard, highlighting the advance recorded in most of the phenological phases in 2017 and 2023. The shortest phenological phase was Flowering (St 6) stage, while the longest was Fruit Development (St 7) stage. The period of low temperatures was necessary to overcome the cooling period in the vineyard studied, which ended in the second half of December month. The thermal requirements to complete berry ripening were around 5,000 heat units (GDD) in the year 2023, which was slightly lower than those required in the year 2017 and 2018. The main conclusion of this study is that high temperatures and drought have reduced the period for the grape harvest in the context of climate change. This is therefore also affecting the other crops and could worsen in the coming years, according to the Intergovernmental Panel on Climate Change (IPCC).

## Acknowledgements

We are thankful to the winery that owns the vineyard, for providing the facilities provided to us for carrying out this research.

## Funding

There was no external funding received for carrying out this research.

## Conflicts of Interest

The authors declare no conflict of interest.

## Notes

### Competing Interest Statement

The authors have declared no competing interest.

